# Social, spatial and temporal segregation in an ant society

**DOI:** 10.1101/002519

**Authors:** Lauren E. Quevillon, Ephraim M. Hanks, Shweta Bansal, David P. Hughes

**Affiliations:** Center for Infectious Disease Dynamics, Penn State University, University Park, Pennsylvania, USA; Department of Biology, Penn State University, University Park, Pennsylvania, USA; Department of Statistics, Penn State University, University Park, Pennsylvania, USA; Department of Biology, Georgetown University, Washington, D.C., USA; Department of Entomology, Penn State University, University Park, Pennsylvania, USA

## Introduction

Sociality can be risky. A chief cost of social living is increased transmission of infectious diseases, due to higher population densities combined with greater contact between susceptible and infected individuals (1,2,3,4). This greater encounter rate has led to a growing interest in the role of social contact structure in infectious disease transmission (5,6,7,8,9,10,11) To capture the dynamics of disease spread within dense groups, epidemiological models are shifting from the principle of mass action, in which infected and susceptible individuals are assumed to mix randomly, to explicitly incorporating patterns of interaction through which infectious agents are transmitted (12). Understanding how interactions impact epidemiology has real-world applications. The growing global connectivity of human communities, coupled with closer proximity to domesticated and wild populations of animals and plants, will impact the incidence of infectious diseases worldwide. Thus, predicting and mitigating the spread of infectious diseases by understanding transmission flow through social contact networks remains a chief One Health imperative (13,14).

Despite the increased propensity for disease transmission inherent to group living, some social organisms have largely overcome this issue. Social insects of the Order Hymenoptera (ants, bees, wasps) are ecologically dominant in almost all terrestrial environments, despite their incredibly dense societies and high degree of genetic relatedness (15). This is not for want of infectious agents-social insects are host to a wide array of pathogens and parasites (16,17,18). Social insects are thought to overcome intense infection pressures through a series of prophylactic and inducible defenses collectively termed “social” or “collective” immunity (19,20). These defenses range from the immunological to the behavioral, including the way colonies are organized and tasks are allocated to workers (21,22,23).

The social and spatial segregation of workers most susceptible to infection is often cited as a major mechanism of disease prophylaxis in social insect colonies (24,25). However, it is unclear if such segregation does indeed occur. We remain unsure because observing individual behavior within a realistic colony has been a formidable task. Here we pursue this avenue of inquiry by testing for the presence of social and spatial segregation in colonies of the carpenter ant, *Camponotus pennsylvanicus*, using analysis of ant social networks combined with individual movement data. *C. pennsylvanicus* is an ant species that has evolved to nest inside dead trees; we mimicked this by maintaining colonies inside wood under complete darkness. We focused on the oral exchange of food, trophollaxis, as the key social interaction of interest because colonies must balance efficient resource flow with mitigating disease spread (26). If social segregation does occur, we would expect to see its signature represented in the trophollactic interactions between castes.

Through the integration of biologically realistic behavioral observations with network and spatial models centered on individual behavior, we ask if ant castes are indeed segregated within the colony. Studies of social insects have greatly benefitted from network analysis because it links local interactions between individuals to the emergent, colony-wide properties that they produce (27,28,29). Several network metrics are of particular relevance to disease transmission, including degree and betweeness centrality. Degree centrality, the number of unique individuals that a given focal ant interacts with, summarizes that individual’s exposure to and potential transmission of infectious agents (11). Although understanding the position of an individual within their social network is important, knowing their spatial context is also crucial for disease flow. Recent advances in automated tracking ability have enhanced our understanding of colony-wide properties such as spatial segregation of different castes (30). However, no empirical studies to date have explicitly linked individual behavior, social network position, and spatial location in a single study. For this reason we combine our network analyses with a statistical analysis of ant movement within the nest.

Finally, we also study the duration and temporal order of trophollaxis interactions because although network and spatial position of individuals are considered important for disease dynamics, the timing of ant-ant interactions is also likely important. We find a number of patterns counter to the strongly prevailing view of social immunity. Within the colony conditions for disease spread would appear ideal. However, by integrating network, spatial and temporal views we find that barriers to disease spread likely exist. It is through this integration of spatial and network analyses with time that might best inform our understanding of disease flow in other complex societies.

## Results

Two colonies of *Camponotus pennsylvanicus*, each containing 75 workers and their queen, were individually housed in an experimental set-up consisting of a wooden nest (area = 63cm^2^) separated from a foraging arena (area = 63cm^2^) by a 4m long maze. Inside the foraging arena, ants had *ad-libitum* access to 20% sucrose solution, water, and a protein source. This 4m separation between the nest and the foraging arena ensured a clear behavioral separation between ants allocated to foraging versus internal nest tasks. Video filming for behavioral analysis was accomplished using a video camera (GoPro Hero2 with modified IR lens, www.ragecameras.com) mounted over the wooden nest illuminated under infrared (IR) light. Ants are unable to perceive light in the IR end of the spectrum and thus its presence was not observed to affect their behavior. Two colonies were filmed over 8 consecutive nights within +/− 30 minutes of 21:00 (when *C. pennsylvanicus* actively forages, L. Quevillon personal observation). An observer watched this playback on a large computer monitor (size) to facilitate behavioral scoring. Each individual ant was observed for trophollaxis events (two ants orally exchanging liquid) for the initial 20 minutes of each recording for each night, leading to a total of over 400 hours of observation (76 ants x 2 colonies x 0.33 hours x 8 nights). The identities of the individuals interacting, the start and stop time of their interaction, and the location of their interaction within the nest was recorded. Additionally, the behavioral identity of the ant during the course of the recording (ie. whether it was an active forager, inactive forager, nest worker, or queen, see methods) was also recorded.

### Static network analysis

There was a significant difference in trophollaxis count between worker types. Foragers engaged in more trophollaxis events than did either nest workers or the queen, although there was no significant difference between active and inactive foragers (post-hoc Tukey HSD on one-way ANOVA, Fig. 1A) However, when the duration of these events is compared across worker types, nest workers had on average the longest trophollaxis exchange, and this was only statistically different from inactive foragers (post-hoc Tukey HSD on one-way ANOVA Fig 1B).

**Figure 1:**
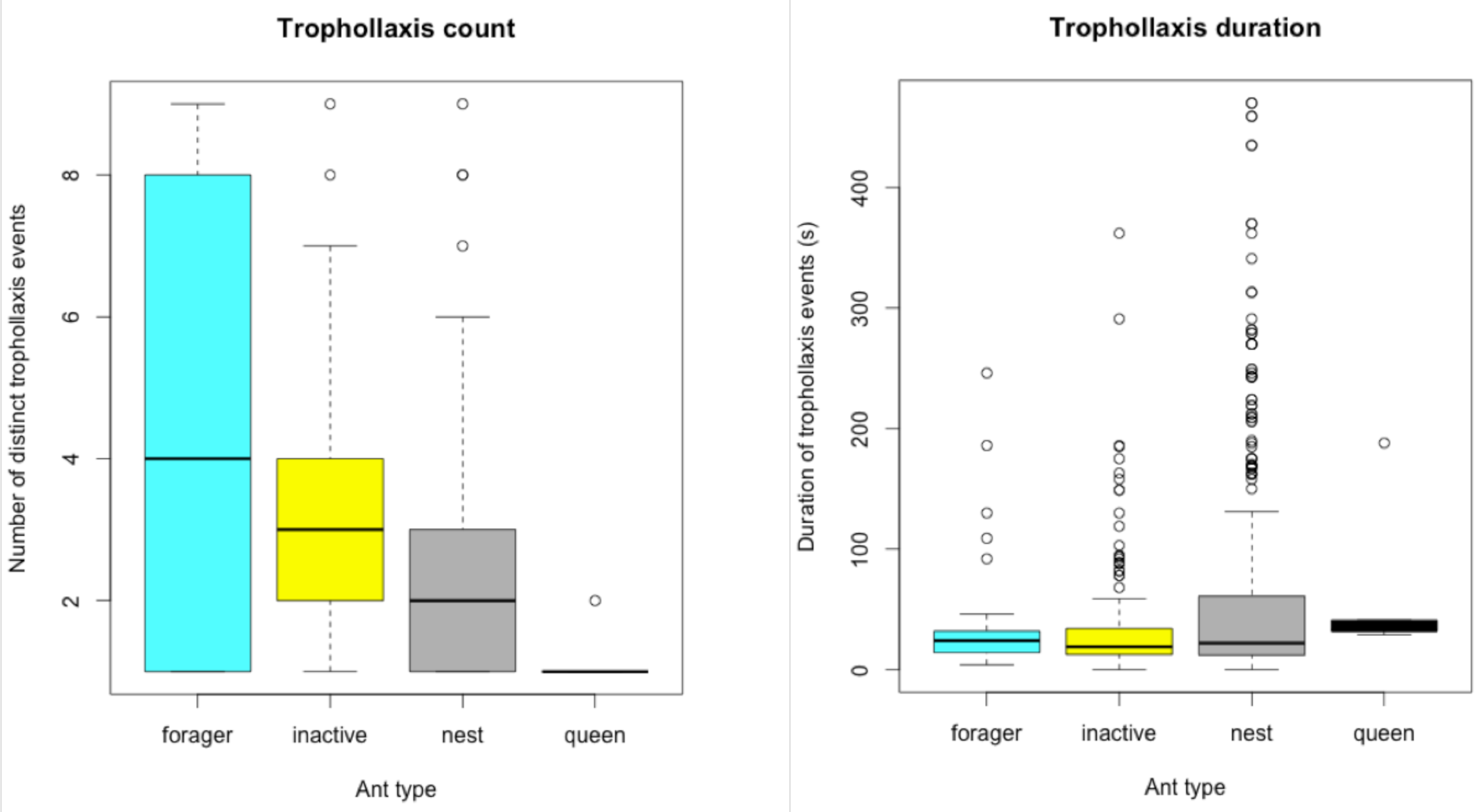
Trophollaxis data for colony 10. a) Trophollaxis count (number of individual trophollaxis events) and b) trophollaxis duration (seconds) as a function of ant worker type. * and ** denote ant worker types that differed significantly in their trophollaxis count or duration (post-hoc Tukey HSD on one-way ANOVA).

**Figure 2:**
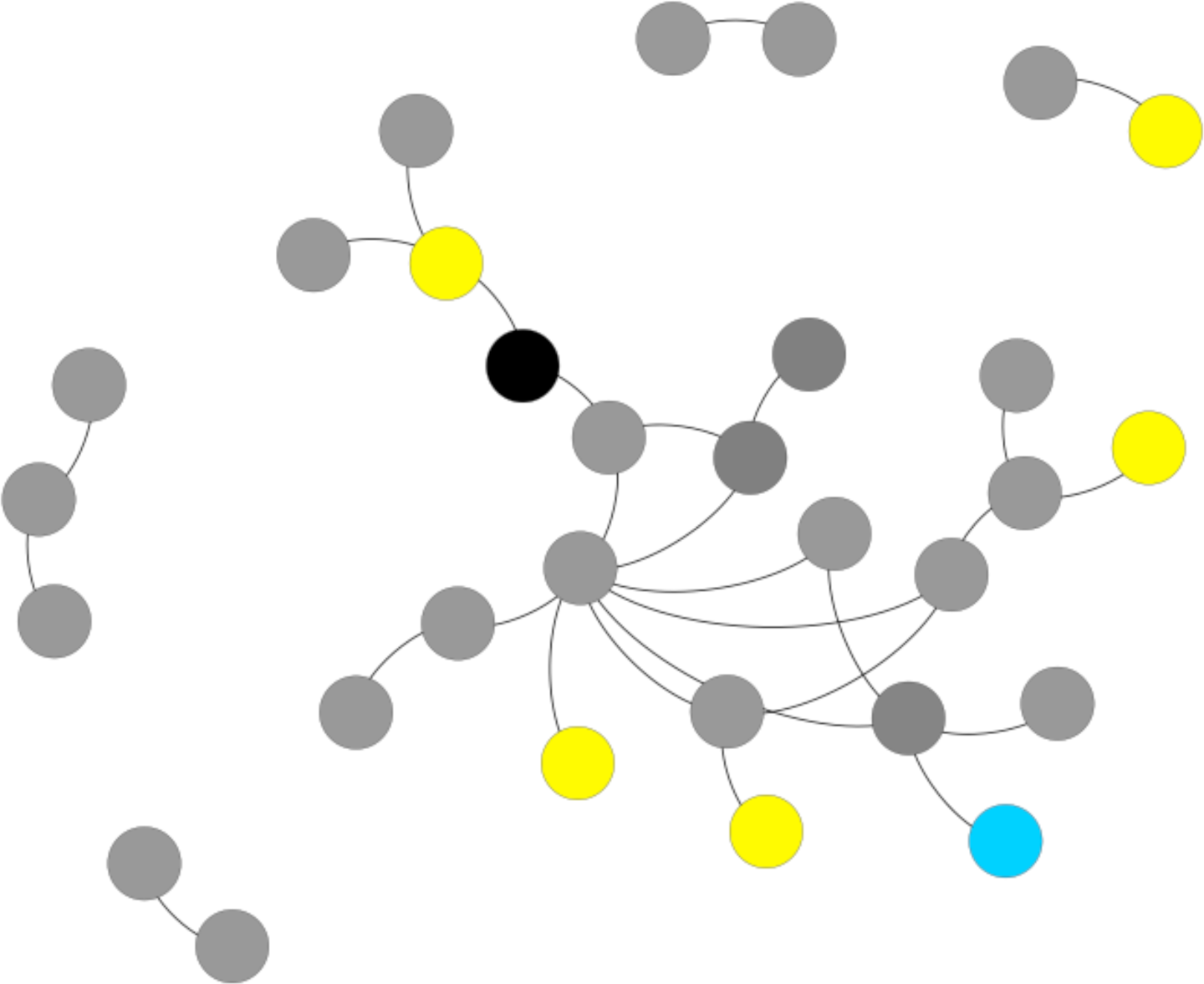
Representative Unweighted Static Network Graph. (Colony 10, June 3rd 2013). Circles represent individual ants; queen (black), nest workers (gray), inactive foragers (yellow), and active foragers (blue). Lines between circles represent trophollaxis interactions between those ants.

Static, unweighted network analyses were conducted on the trophollaxis interaction for a single colony using the package ‘iGraph’ (31) implemented in R (32). Active foragers (ants who were observed to forage during the video recording) had a higher degree (number of unique individuals with which they engaged with through trophollaxis) than inactive foragers, nest workers, or the queen. This represents an average of 2 additional unique individuals that foragers exchanged food with compared to the queen. While the queen had an average degree of 1, the identity of the individual she interacted with was not consistent across nights.

### Ant movement and spatial analysis

For each colony, individual ant movement patterns were investigated by randomly choosing five known foraging ants, five known nest workers, and the queen to have their spatial movement data recorded. The wooden nest in which ants were housed was gridded to a resolution of 1cm^2^, and the cell locations where the majority of the ant’s body was located as well as the time stamp when it was in that location were recorded for the entire 20-minute duration of the video for each of the 8 nights. The residence time spent in each cell was recorded and summed over all ants to determine nest spatial use. In both colonies, the queen Residence times in each cell and transitions to neighboring cells were used to fit a continuous-time discrete-space random walk model for ant movement behavior, where were used to calculate a movement or transition rate between cells.

The average spatial usage of foragers, nest workers, and the queen is given in Fig. 3. Foragers occupied a greater proportion of the nest than did either non-foraging nest workers or the queen. The queen was largely immobile in both colonies, though in one colony (Col10), the queen spent some time in 3 of the 4 chambers of the nest.

**Figure 3:**
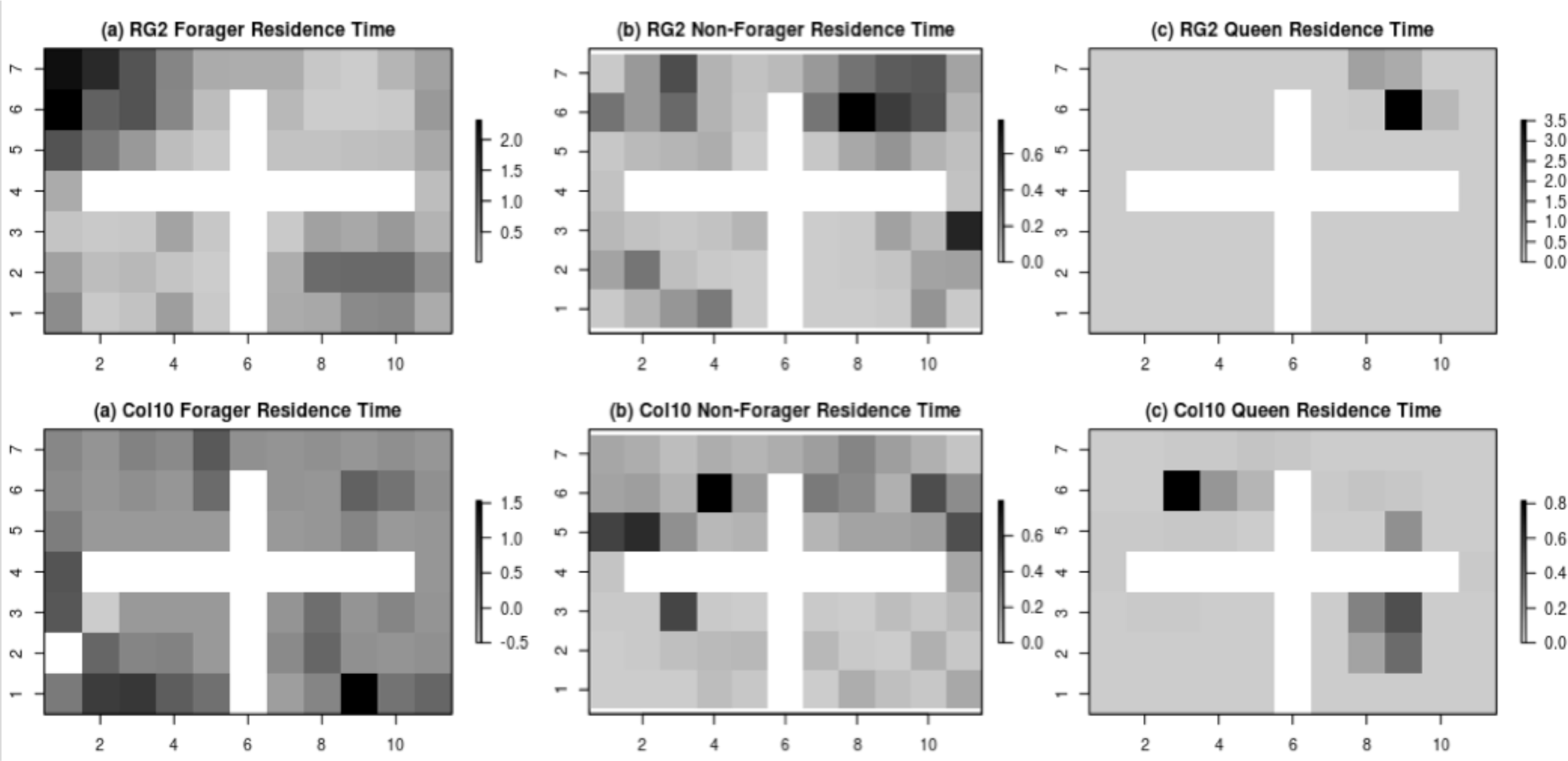
Segregated Use of Nest Space. Aggregated residence times in ant-days for queens, active foragers, and non-foraging ants from two colonies (RG2 and Colony 10).

To test for differences in movement behavior, we used a continuous-time discrete-space Markov chain model for ant movement (33) that allows for testing differences in movement behavior between worker types in response to spatial covariates. We tested for differences in overall mean movement rates between foraging and non-foraging nest workers, and for changes in movement rates when in the same chamber as the queen. We also tested for directional bias in movement behavior toward or away from the queen (i.e., queen avoidance). Results of this analysis show that in both colonies non-foraging ants are more mobile (have higher movement rates) than are foraging ants while in the nest (p<10＾-10, T-test). There was no evidence of directional queen avoidance by foraging or non-foraging ants in either colony, but there was strong evidence in one colony (RG2) that foraging ants move faster when near the queen then when in another chamber (p<10＾-14, T-test) and that non-foraging ants tend to move slower when in the same chamber as the queen (p<0.01, T-test). In the second colony (Col10), non-foraging ants also tend to move slower when near the queen (p<.01, T-test), but there is small evidence that foraging ants move faster near the queen (p<.3, T-test). This discrepancy is likely due to the increased movement of the queen in the second colony, which obscures the spatial movement signal.

### Time-ordered (time-dependent) social network analysis

Social network data has traditionally been analyzed as a time-aggregated or static graph, in which the timing of interactions and their order is ignored. However, this timing and order is crucially important for dynamic flow processes, such as disease transfer (34). We re-analyzed the ant interaction data using the package ‘timeordered’ (35) implemented in R. This specifically incorporates the time stamp of interactions when computing network metrics, and allows for a much more biologically meaningful picture of intra-colony interactions of import to disease. Fig. 4 shows a representative time-ordered network graph. Based on the timing of interactions, returning foragers were never actually observed to interact in a way necessary for disease transmission.

**Figure 4:**
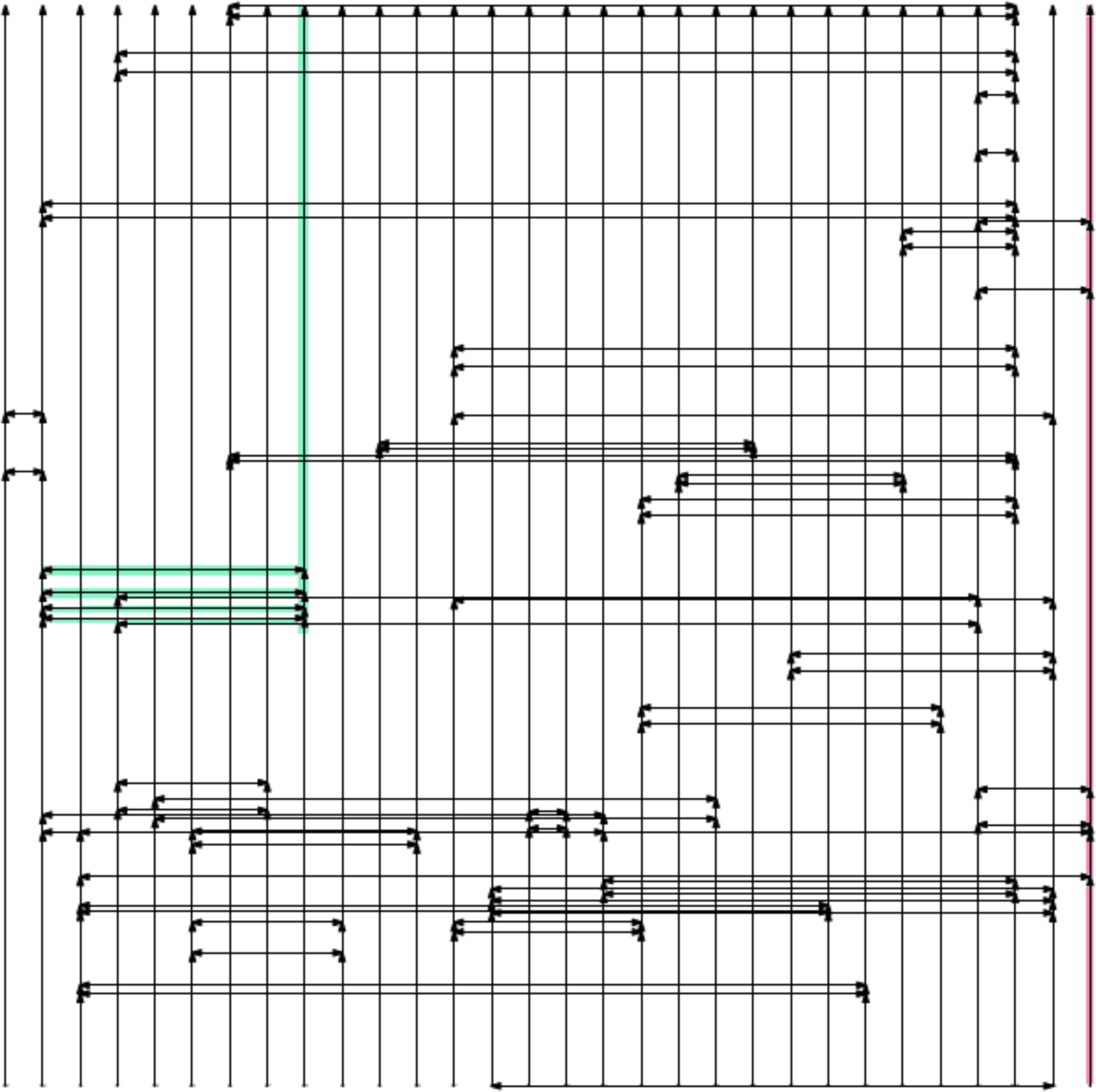
Representative Time-Aggregated Network Graph. (colony 10, June 10th, 2013). Each vertical line represents an indivdual ant, time increases up the vertical axis. Horizontal lines represent trophollaxis events between the indivudal lines that they connect. The queen is highlighted in red, and a foraging ant that has returned to the nest is highlighted in green. Note that the queen and foraging ant never interact in a temporally meaningful way, despite their overall connectivity within a static network representation.

## Discussion

The results of this study provide a comprehensive view of social, spatial and temporal segregation of different ant types within the colony. Static network analyses revealed that actively foraging ants engage in social food exchanges with more individuals than either nest workers or the queen. This is a surprising finding given that actively foraging ants have the highest disease exposure of all ants, and thus social immunity theory would predict that their contact with susceptible nest mates should be minimized (17,19). When the duration of trophollaxis events is taken into account, however, there are not statistically significant differences between foragers and nest workers. This could be a function of the biological limits to oral food transfer in *C. pennsylvanicus* and is worth further investigation. This component of the trophollaxis data is in accordance with what social immunity theory would predict (ie. foragers don’t engage long, and therefore dangerous, trophollaxis interactions with nest mates).

In addition to social position of ants within the colony, we were also interested in the spatial activity of such ants. Analysis of nest spatial usage showed that foragers are spatially promiscuous, nest workers are less so, and the queen hardly moves. While the queen’s lack of movement syncs well with our predictions from social immunity studies, the expansive movement of the foragers is counterintuitive; theory predicts that foragers should be avoiding internal areas of the nest. However, in one colony (RG2), it does appear that foragers may be modulating their speed in response to their social environment. When foraging ants were in the same chamber as the queen, they moved faster than their nest worker counterparts. By moving faster near the most important individuals in the colony, foragers may be reducing the potential transmission of any infectious agents that they may have been exposed to.

The static network analyses of colony social organization and the spatial promiscuity of foragers to queens reveal an ant society not particularly well suited to the prevention of disease transmission. However, the inclusion of temporal data makes this situation far less dire than what it appears. When the timing and order of trophollaxis interactions are taken into account, foragers and the queen never interact in a way that could lead to the biologically meaningful transfer of disease (ie. after a forager has come back into the nest after a foraging trip, carrying some pathogen that might transfer to the queen via close proximity or oral food exchange). Thus, the timing of social interactions coupled with movement rates provides evidence for behavioral prophylaxis within *C. pennsylvanicus* colonies.

Through the incorporation of social interactions, individual movement data, and the timing of social interactions, we now have a better understanding of how disease prophylaxis could be accomplished in *C. pennsylvanicus* ant societies. Had the timing of interactions and movement been ignored, a different picture invalidating tenets of social immunity theory would have emerged. This provides further evidence for the growing argument that temporal information and meaningful behavioral interactions should be included into social network analyses if we are to make biologically accurate conclusions (34). Laboratory studies involving animal behavior benefit from the incorporation of environmental complexity and ecological realism. We encourage the continued advancement of experimental set-ups if we are to gain a true understanding of how social insect societies are structured.

Having provided a necessary null model of colony organization in the absence of disease, future experiments in which laboratory infections are combined with network analyses will further inform the extent to which colony organization reduces disease transmission in social insect societies. Such studies will also afford us the ability to synchronize theoretical predictions from agent-based modeling approaches (36) with empirical data that will allow for enhanced model parameterization. Social insect societies are a powerful model system for investigating how perturbations in social structure can influence disease transmission dynamics. However, to realize their full potential we advocate for continued inquiry through the use of biologically meaningful behavioral interactions that include temporal information.

## Methods

### Ant colony set-up and filming

Two queen-right *Camponotus pennsylvanicus* colonies were collected from field sites in Pennsylvania, U.S.A. in December 2012. Seventy-five worker ants were haphazardly selected from each colony and were individually labeled. Labels consisted of numbers printed on photo paper that were affixed to the ants’ gasters with optically clear nail polish. The labeling was not observed to alter the ants’ behavior or interactions (L. Quevillon, personal observation).

The labeled ants and the queen were housed in a nest set-up consisting of a four-chambered wooden nest (total area = 63 cm^2^) that was gridded to a resolution of 1cm^2^ and covered with a plexiglas top. This nest was contained within a filming box so that nest conditions were always dark. The nest was separated from a sand-bottomed foraging arena (total area = 63 cm^2^) by a 4 m long maze. The length of the maze ensured that there was a clear separation between workers allocated to foraging versus internal colony tasks (L. Quevillon, personal observation). Inside the foraging arena, ants had *ad libitum* access to water, 20% sucrose solution and mealworms.

Each colony was filmed at +/− 30 minutes of 21:00 on 8 consecutive nights in June 2013 using a GoPro Hero2 camera with a modified IR filter (RageCams.com) illuminated under infrared light (Canon CMOS IR light). Infrared light, which ants are unable to perceive (reference), was not observed to affect ant behavior.

### Video analysis and ant worker classification

For each night of filming, the trophollactic interactions of every ant inside the nest were individually observed. Due to degradation of IR light intensity while filming, only the first 20 minutes of each video were analyzed. For each trophollactic interaction that was observed, the ant identities, start time, stop time, and location within the nest were recorded. Additionally, the overall behavioral category of each ant on each day was recorded (i.e. nest worker, forager, non-active forager, queen). Nest workers were ants that were never observed to leave the nest, foragers were ants that actively left the nest during the course of the video segment, and inactive foragers were ants that had been witnessed to leave the nest in video segments on previous days, but which did not leave the nest during the video segment being currently analyzed.

### Trophollaxis count and duration

The number of trophollaxis events and their duration for each individual in Colony 10 was recorded as given above. To test for differences in both trophollaxis count and duration as a function of ant type (ie. forager, inactive forager, nest worker, or queen), a one-way analysis of variance was conducted using .aov in R. Post-hoc tests for differences (Tukey HSD) were then used to determine which ant types had significant differences from each other.

### Static network analysis

Network metrics were analyzed for colony 10 for each night of observation. Unweighted, static network analyses were conducted using the iGraph package (31) implemented in R (32). Metrics analyzed for each individual ant included degree, betweeness centrality, closeness centrality, and Burt’s constraint.

### Spatial movement analysis

The time-referenced spatial locations of the queen, 5 forager ants, and 5 randomly chosen nest-worker ants were recorded for each night. We used a continuous-time discrete-space agent-based random walk model (33,37) to make inference about ant movement behavior. The CTDS framework is notable in that it allows for inference on both directional (e.g., queen avoidance) and location-based (e.g., variable movement rates in different nest chambers) movement mechanisms. Additionally, Hanks et al., (2013) have shown how inference can be made on CTDS movement models under a standard generalized linear modeling (GLM) framework, which leads to intuitive inference and efficient computation. Drawing on standard continuous-time Markov chain models (e.g., 38), if an ant is in cell *i* at time *t*, then define the rate of transition from cell *i* to a neighboring cell *j* as *λ(ij)*. The total rate *λ(i)* at which ants move (transition) out of cell *i* is the sum of the rates to all neighboring cells: *λij* = *λij*, and when the ant moves, the probability of moving to cell *k* (instead of to another neighboring cell) is the ratio: *λikλi*.

To model ant movement behavior near the queen, we will model *λ(ij)* as a function of a spatial covariate which measures the distance from the queen’s most used locations (DFQ) at each grid cell (Figure 3). To examine local behavior, the DFQ covariate was set to be constant out of the queen’s chamber. The DFQ covariate is location-based and will allow us to model differences in movement rates when near or far from the queen. We also considered a directional covariate, a gradient of the DFQ covariate (GDFQ). The GDFQ gradient is a directional vector that points towards the queen, or along the direction of steepest ascent of the DFQ covariate, and the GDFQ covariate will be different for the transition rates to neighboring cells in different directions, thus allowing for directional preference in ant movement. We also consider potential differences in movement behavior between foraging (F) and non-foraging (NF) ants, with F=1 for foraging ants and F=0 otherwise, and NF=0 for foraging ants and NF=0 otherwise. We model the movement rate *λ_k_(ij)* of the *k*-th ant from cell *i* to cell *j* as a function of interactions of these covariates and corresponding regression parameters {β}:

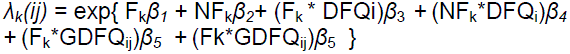

Differences in overall movement rates between foragers and non-foragers will be represented by differences in *β*_1_ and *β*_2_, with positive values corresponding to higher movement rates. Positive values of *β*_3_ correspond to higher movement rates of foraging ants when far from the queen, and decreased movement rates near the queen. Positive values of *β*_5_ correspond to preferential directional movement by foragers away from the queen (in the direction of the increase in the gradient of DFQ). The parameters *β*_4_ and *β*_6_ correspond to the response of non-foraging ants to DFQ and GDFQ, respectively. Hanks et al. (2013) have shown that inference on the parameters in this movement model can be accomplished using a Poisson GLM, which we fit using the ‘glm’ command in R. Results are summarized in Table 1.

**Table 1:**
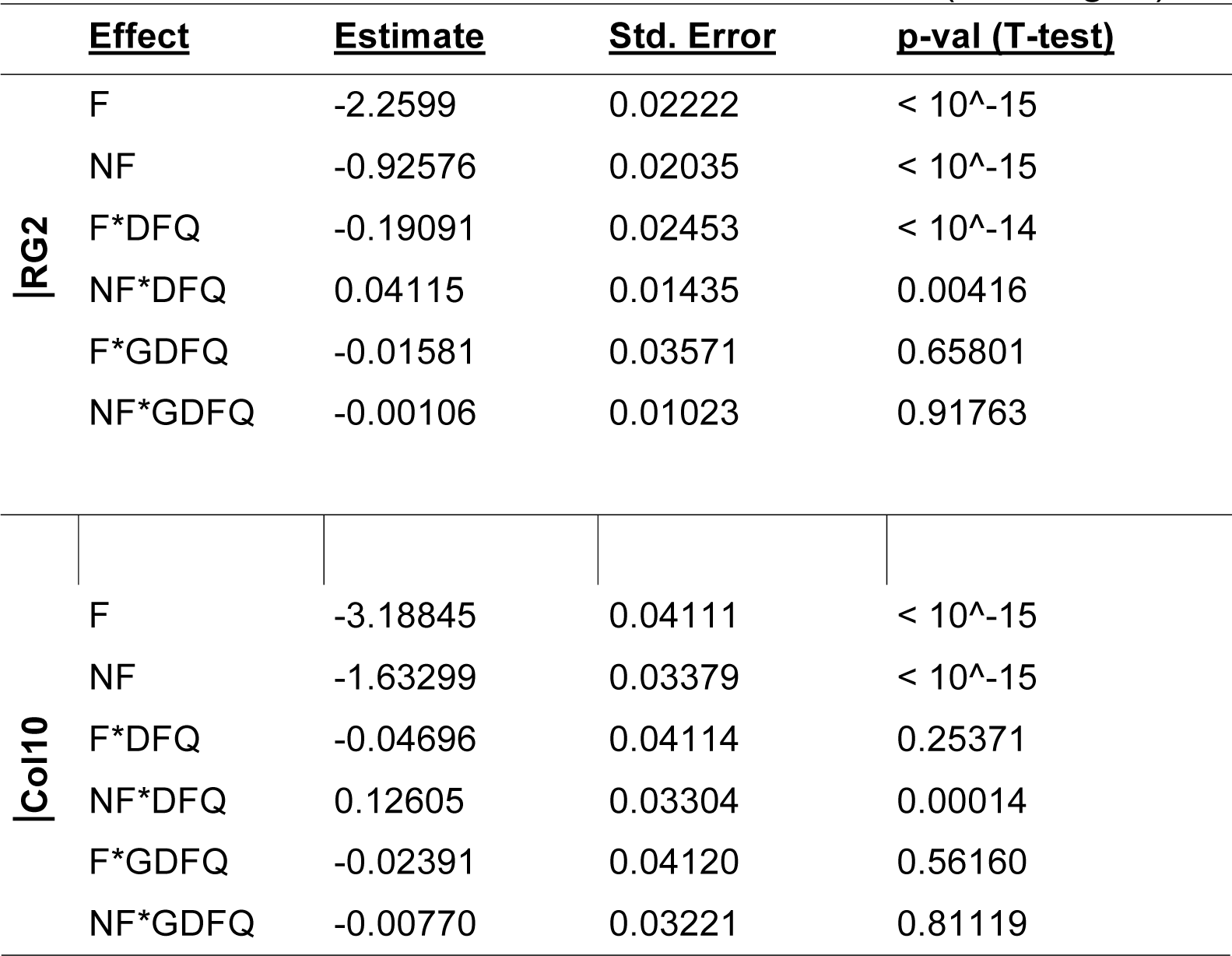
Inference on ant movement parameters in a continuous-time random walk model of ant movement in two ant colonies (See Fig. 3)

### Temporal (time-ordered) network analysis

Interactions from the static network analyses were re-analyzed including the time-stamp of when the interactions occurred. Temporal networks were constructed using the package ‘timeordered’ in R. The time to interaction between foraging ants and the queen was calculated using the function ‘shortesttimepath’.

